# Computational Modelling of Immune Interaction and Epidermal Homeostasis in Psoriasis

**DOI:** 10.1101/2023.02.23.529657

**Authors:** Dinika Paramalingam, Bowen Li, Nick J. Reynolds, Paolo Zuliani

## Abstract

Psoriasis is an incurable chronic inflammatory skin disease characterised by immune cytokine-stimulated epidermal hyperproliferation. This results in the skin becoming red with scaly plaques that can appear anywhere on the body, decreasing the quality of life for patients. Previous modelling studies of psoriasis have been limited to 2D models and lacked cell-cell interactions. We have developed a 3D agent-based model of epidermal cell dynamics to gain insights into how immune cytokine stimuli induces hyperproliferation in psoriasis to better understand disease formation and structural changes. Three main keratinocytes, stem, transit-amplifying (TA), differentiated and T cells, are modelled with proliferation and division governed by various nutrients and immune cytokines. Each cell has a set of attributes (growth rate, division probability, position, etc) whose values are governed by processes such as monod-based cellular growth model, probability-based division based on calcium and cytokine concentration and various forces to form the epidermal layers. The model has 2 steady states, healthy (non-lesional) and psoriatic (lesional) skin. Transition from healthy to psoriatic state is triggered by a temporary cytokine stimulus which causes hyperproliferation to occur, a hallmark of psoriasis. This results in the deepening of rete ridges and thickening of the epidermal structure. Model outputs has been validated against population ratios of stem, TA, differentiated, and T cells, cell cycle and turnover times in vivo. The model simulates the structural properties of epidermis, including layer stratification, formation of wave-like rete ridges, change in epidermal height and length of rete ridges from healthy to psoriasis. This has provided some insights on the complex spatio-temporal changes when transitioning between the 2 steady states and how a shot of temporary cytokine stimulus can induce different severity of psoriasis and alters proliferation between healthy and psoriatic skin in line with known literature. This provides the basis to study different cytokine simulation variations of psoriasis development and tracking of cell proliferation in the lab. It also provides a baseline to model the effects of psoriasis treatments such as narrowband-ultraviolet B (NB-UVB) or biologics and predict potential treatment outcomes for patients.

## Introduction

Psoriasis is a chronic and disabling inflammatory skin disease that affects about 25 million people in North America and Europe [1], and approximately 2-3% of the UK population, affecting both genders equally. The disease occurs at any age and reduces the quality of life [2]. There is no cure for psoriasis, however, it is a treatable condition with various types of treatment options available depending on the severity of the disease ranging from topical treatment, such as topical corticosteroids to systemic therapies, such as methotrexate, to phototherapy treatments, such as Narrow-band Ultraviolet B (UVB)[2], [3]. The mechanism of action for each type of treatment varies, and one of the most commonly used treatment is phototherapy, UVB, in particular for moderate to severe psoriasis [3, 4, 5, 6]. UVB works by various mechanism of actions such as altering the cytokine profile, apoptosis of keratinocytes for clearance, promotion of immunosuppression and other mechanisms like cell-cycle arrest [4].

The development of psoriasis is complex and multi-factorial and can be triggered by an abnormal immune response such as an infection that causes T cells to secrete cytokines to drive hyperproliferation in the epidermis. This causes keratinocytes to contain a higher level of proliferation as compared to normal keratinocytes [7]. Therefore, giving rise to a proportionally higher cell population in psoriatic epidermis. This not only causes a thicker epidermis but gives rise to deeper rete ridges and an absent granular layer [8, 9].

There are various cytokines secreted by T cells such as Interleukin (IL) 17, IL-22, Tumour Necrosis Factor *α* (TNF-*α*) which affect the proliferative cells to grow faster and divide more quickly causing a significantly shorter turnover time. In response to this, proliferative cells increase their production of endogenous growth factors (GF), which in turn results in this positive feedback loop for it to maintain its psoriatic state (Figure 1).

**Figure 1:**
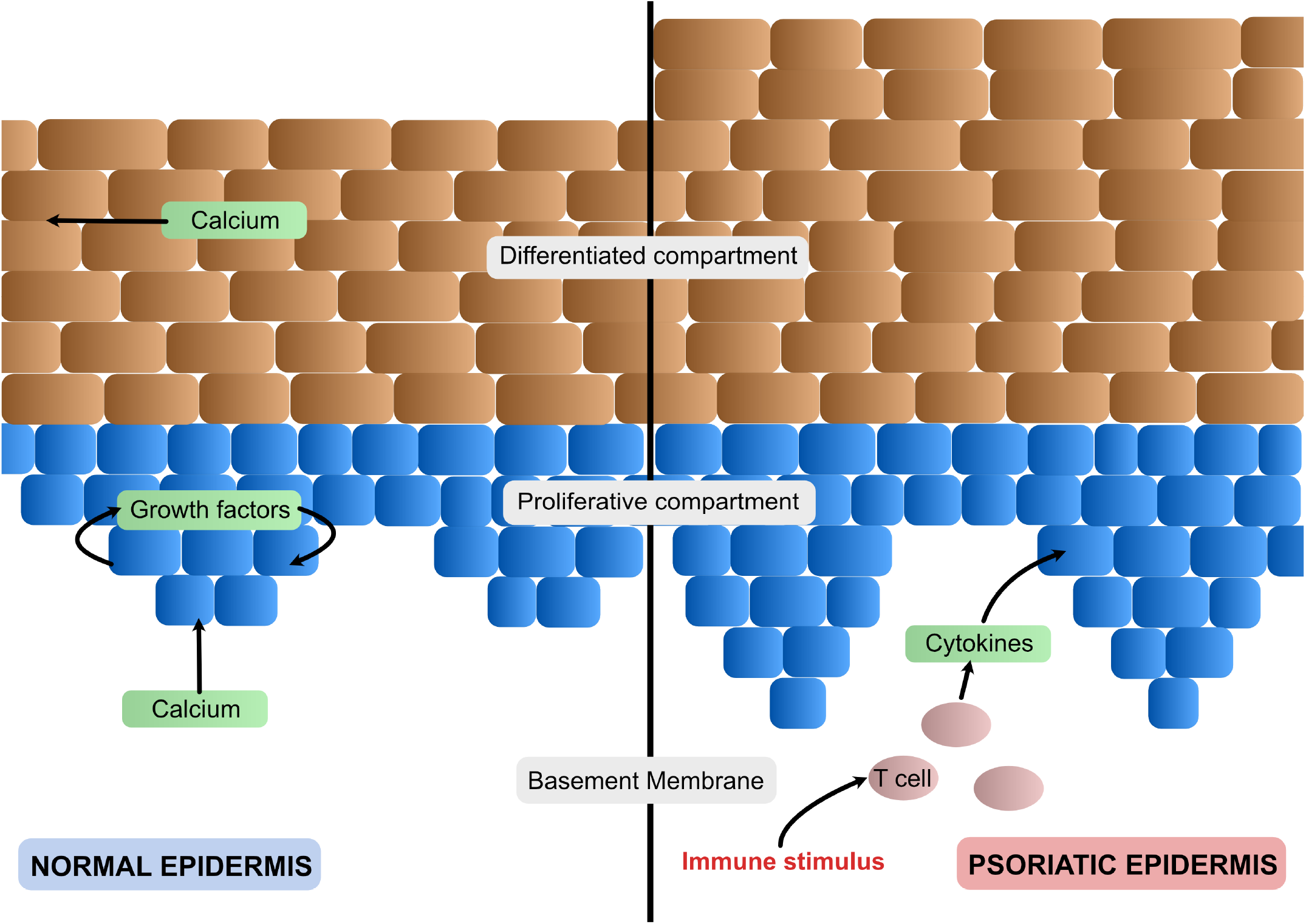
Schematic of healthy (left) and psoriatic (right) epidermis. The epidermis is made up of two main compartments - proliferative and differentiated. The proliferative compartment is made up of stem and TA cells while the differentiated compartment is made up of terminally differentiated cells. In the healthy epidermis, the rete pegs are more swallow and the overall epidermal height is shorter than in the psoriatic state. The nutrients involved in its development can vary. For example, in this case, calcium from the dermis aids proliferation and endogenous growth factors are produced by the stem and TA cells which also regulates its growth. In psoriasis, an immune stimulus triggers T cells to produce cytokines such as IL-22 and TNF*α*, which are diffused into the epidermis and drive stem and TA cells into hyperproliferation. As a result, more cells are produced causing the rete pegs to deepen and the overall epidermis height to increase.

**Figure 2:**
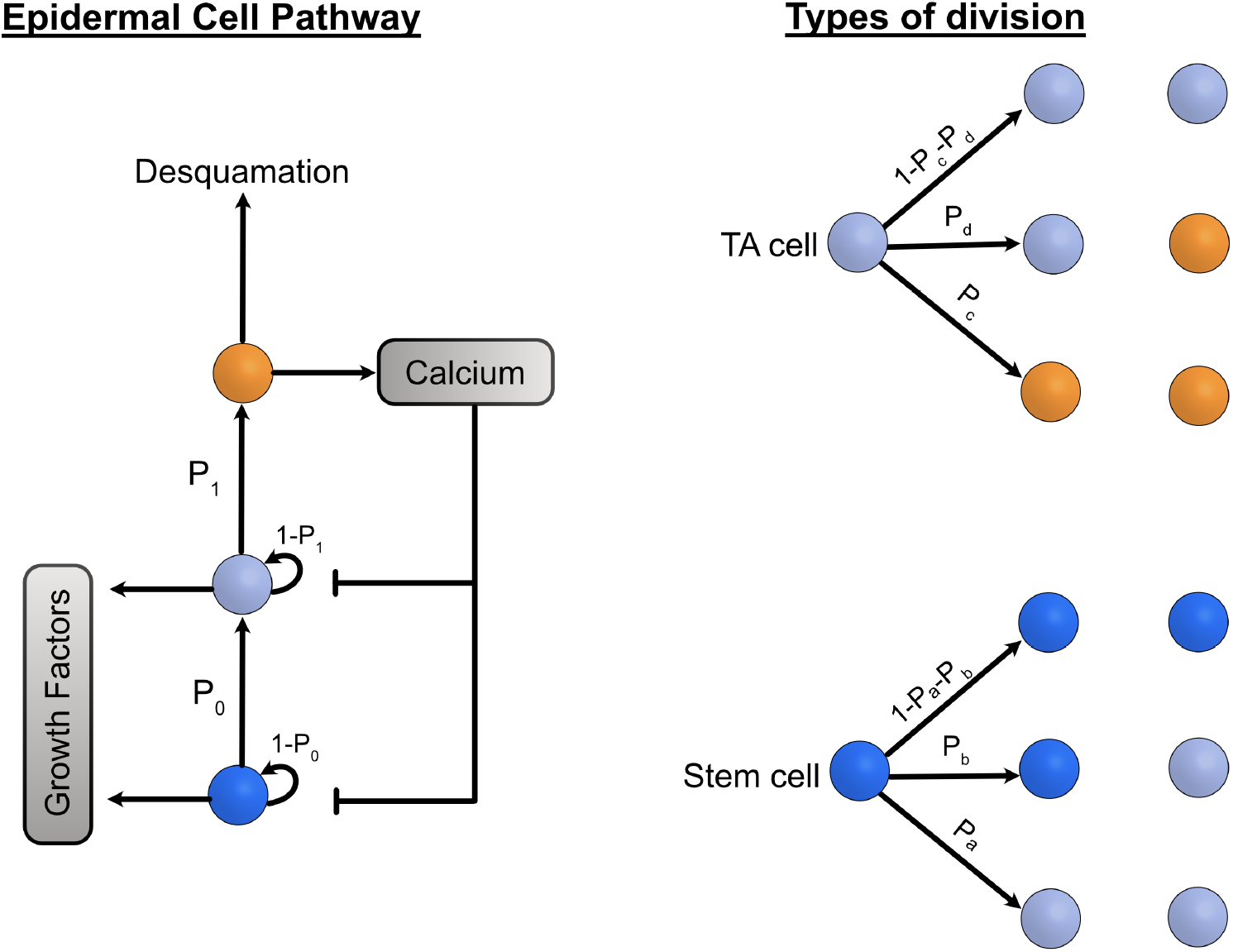
(Left) Description of the epidermal cell pathway from a stem cell (in dark blue). Stem and TA cells growth and divide to either one of the three division methods where their probabilities to have a daughter cell of a different type (i.e. TA or differentiated cell, respectively) is *P* 0 and *P* 1 while having a daughter of the same type is 1 - *P* 0. Stem and TA cells produce growth factors that drive their growth and proliferation and differentiated cells secrete calcium which inhibits stem and TA cells growth in order to maintain a steady state. (Right) Stem and TA cell division can be divided further to the three different ways and are based on the probabilities *Pa, Pb* for stem cells and *Pc, Pd* for TA cells.

We present a 3D computational model to simulate epidermal development and homeostasis and its transition to psoriasis using immune cytokine stimulus. The development of the model aims to provide an attractive solution to some of the challenges and limitations in previously developed computational models such as:

- Modelling the cell-cell interactions between keratinocytes and T cells
- Mimicking the wave-like structure of the basement membrane as seen in vivo
- Introducing a single immune cytokine stimulus to trigger psoriasis
- Model maintains two steady states - healthy (non-lesional) and psoriasis (lesional)

## Methodology

The model consists of three main sub-processes - biological, physical and chemical processes. The biological processes focus on how the different cell types grow and divide, while the physical processes focus on the spatial aspects such as rete peg formation, mechanical relaxation, spatial regulation for cell division on the wave-like basement membrane and a flexible basement membrane when transitioning to psoriasis. Finally, the chemical processes focus on how the nutrients, endogenous GF, calcium, and immune cytokine are regulated in the model.

### Biological Processes

The biological processes consists of cell growth and division which are based on the nutrients and cytokines involved and consist of two steady states, healthy (non-lesional) and psoriatic (lesional) epidermis.

### Cell growth

The growth model for both healthy and psoriatic epidermis are based on various nutrients and immune cytokines that have been found in literature and in clinical studies [7, 10, 11]. Cell division of proliferative stem and TA cells are based on having two interconvertible modes as mentioned in [12] where proliferative cells change their division type based on an expanding or balanced state. (Figure 3).

**Figure 3:**
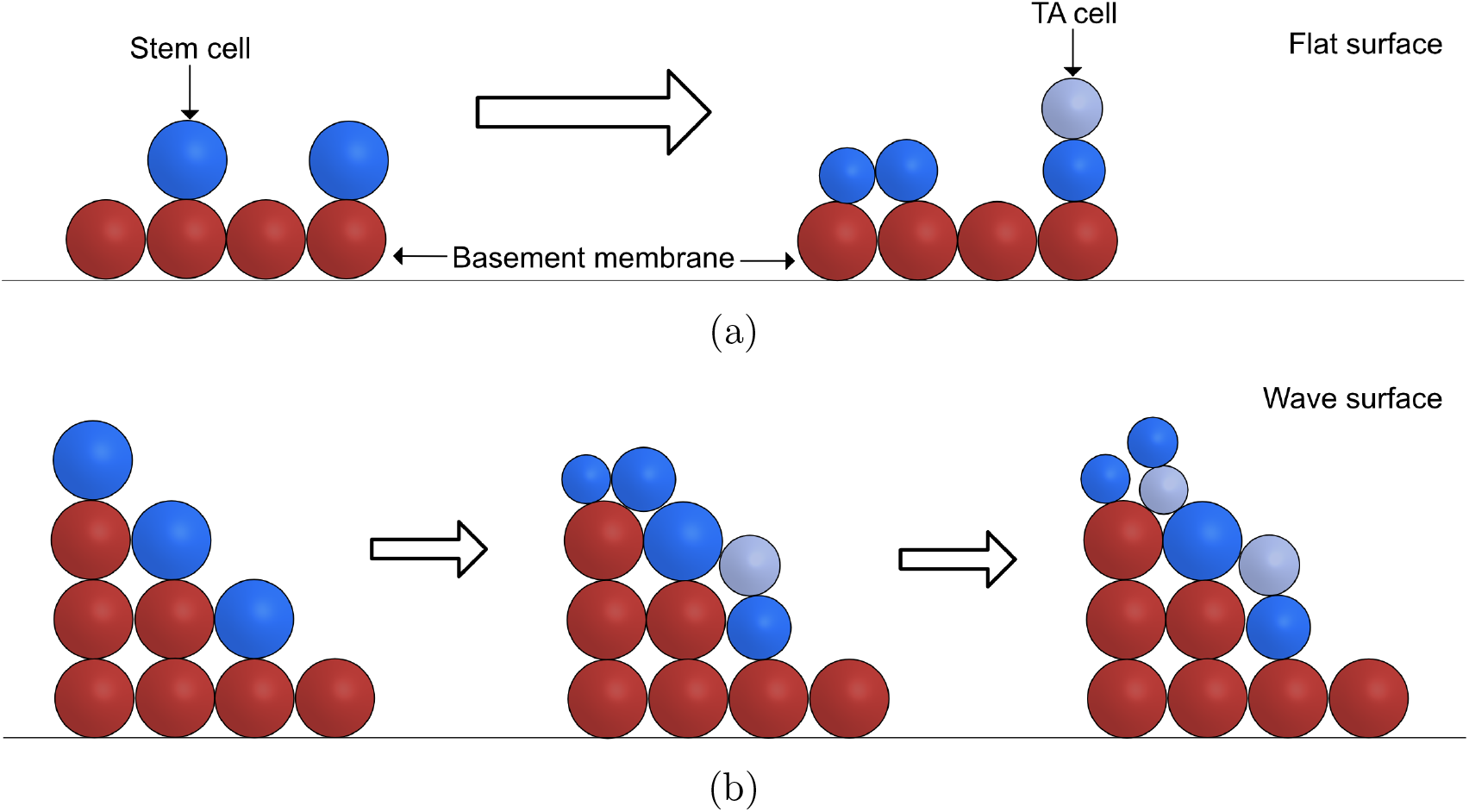
(a) Division direction for stem cells. If the stem cell undergoes self proliferation, it will divide horizontally while in asymmetric division, the TA daughter cell will be placed on top (i.e. vertically). This is straightforward if the basement membrane was a flat surface. (b) In the wave surface, a stem cell divides asymmetrically, a TA cell could be positioned on top of the basement membrane. However, there will be an issue if another stem cell around it divides to produce a daughter stem cell. The daughter stem cell, will end up positioned on top of the TA cell instead.

As each cell grows, its mass increases by consuming the nutrients in its environment. The growth process is described by the following equation:

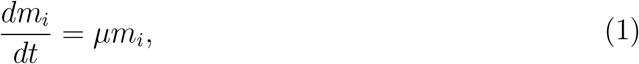

where *µ*_*i*_ is the specified growth rate and *m*_*i*_ is the biomass of the *i*th cell. The growth rate *µ*_*i*_ is determined by the following growth equations:

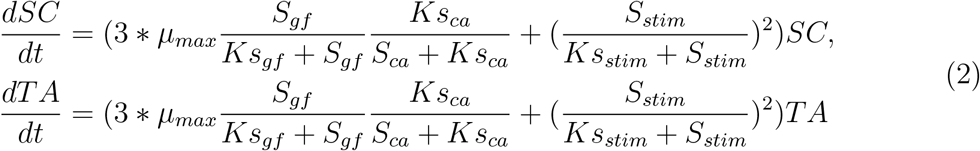

where *µ*_*max*_ is the maximum growth rate specified, *S*_*gf*_ and *S*_*ca*_ are the current concentration of endogenous GF and calcium within the voxel respectively, and *Ks*_*gf*_ and *Ks*_*ca*_ are the half-velocity constants for growth factor and calcium. *SC* and *TA* are the number of cells in the system. Immune cytokines stimulus, *stim*, is added into the growth equation to trigger hyperproliferation when transitioning to psoriasis. *S*_*stim*_ and *Ks*_*stim*_ represents the current concentration and half-velocity constant for immune cytokine stimulus, respectively.

Cellular growth based on the Monod kinetics is driven by the nutrients in the voxel where each cell resides. The nutrients present in each voxel are growth factors and calcium in both the healthy and psoriatic epidermis. In psoriasis, *stim* is the additional nutrient. Haldane inhibition is based on the calcium concentration in each voxel and aids in the maintenance of the epidermal structure. Differentiated cells secrete calcium as they migrate upwards towards the stratum granulosum layer before desquamating. The calcium concentration in the system increases and the proliferating cells will “sense” this high concentration to slow down their growth, which in turn, maintains a steady state.

### Cell division and desquamation

Cell division occurs as a result of cell biomass growth and desquamation occurs to emulate the shedding of differentiated cells at the top of the epidermis. Division occurs if the diameter of a cell reaches the specified threshold value and the cell splits into two daughter cells. The mass of one daughter cell is a randomly assigned value between 40-60% of the parent’s total mass and the other daughter cell receives the rest. In addition, one daughter cell will take the position of the parent’s while the other daughter is placed randomly in either a horizontal or vertical direction from the parent cell depending on its type. Stem cells are known to be only found along the basement membrane and so, if the daughter cell is a stem cell, it will be placed adjacent to its parent while a TA cell can be placed in either direction [13, 14].

In addition, calcium is used to control the division probabilities in the system. The cell division probabilities are determined by the state the model is in. In the expanding stage, where the simulation starts with just stem cells, self proliferation occurs at the faster rate and once the model reaches a steady state, the model switches to a balanced stage to maintain the cell density at a steady state. The following equation describes the division equations used for stem (*Pa* and *Pb*) and TA (*Pc* and *Pd*) cells.

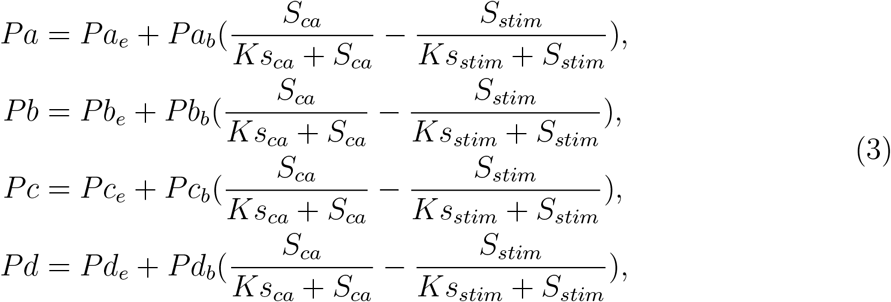

where *Pa* and *Pc* represent symmetric division and *Pb* and *Pd* represent asymmetric division while the remainder is self proliferation. The first variable (i.e. *Pa*_*e*_, *Pb*_*e*_, *Pc*_*e*_, *Pd*_*e*_) in the equation is the probability used during the expanding state to get the model to a steady state and switches to a balanced state with a different set of probabilities (i.e. *Pa*_*b*_, *Pb*_*b*_, *Pc*_*b*_, *Pd*_*b*_) governed by a Hill function based on the calcium concentration in the system. During the transition to psoriasis, immune stimulus, *stim*, causes the model to go back into an expanding state resulting in hyperproliferation (see Figure 1).

TA cells are known to divide a finite number of times (4-5 times) before terminally differentiating [7, 15, 16]. An additional rule has been added to TA division, where the maximum number of times a TA cell can divide is 4 in healthy epidermal development and 5 in psoriasis, based on a previous model developed [7].

### Endogenous GF and Calcium production

The Monod-based kinetics allows stem and TA cells to produce GF and differentiated cells to secrete calcium into the environment. The concept of GF production by stem and TA cells is based on a previously developed ODE model [10], while calcium is stored in the proliferative cells as they grow, divide and terminally differentiate and gets secreted when a differentiated cell reaches the stratum granulosum layer.

### Physical Processes

The model consists of various physical processes to solve issues related to overlapping cells during growth and division, development of the rete pegs and the transition from healthy to psoriatic state where there is morphological change to the shape and height of the rete pegs. In addition, different cell types have different cortical and adhesive forces [17, 18].

### Formation of rete pegs and stem cell initialisation

The structure of the basement membrane is known to be of a wave-like structure instead of a flat surface. To emulate this morphology, the formation of the rete pegs is assumed to be of a sine wave structure and made out of spherical agents holding the structure. The LAMMPS command “create atoms” allows us to specify the x-,y-, and z-axis for atoms to be created based on region blocks and the wave was created using the following formula:

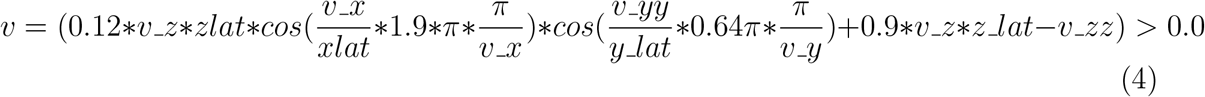

where the variable *v* stores the formula to create the waves. The first part of the equation, 0.12 *v*_*z zlat*, affects the depth of the wave while *cos*(*v*_*xx/xlat* 1.9 *π π/v*_*x*) affects the width of the wave on the *x*-axis. The second part, *cos*(*v*_*yy/ylat* 0.64 *π π/v*_*y*), controls width of the wave on the *y*-axis. Lastly, 0.9 *v*_*z zlat v*_*zz*, affects the top peaks of the waves. Having a larger value (i.e. more than 0.9), results in a flatter top.

As the basement membrane surface was no longer flat, stem cell initialisation had to be modified and a new fix was implemented to ensure that the stem cells were able to initialise on top of the basement membrane rather than having some cells initialised in the top peaks. A “fix” is a type of operation command in LAMMPS that can have multiple functions such as applying constraints to atoms, creating boundaries and so on and so forth. The fix is written in C++ and is part of the software’s source code. This makes it cleaner and easier for the user to specify parameters in the input script rather than in the source code itself. The new fix *fix_pso_atoms* was implemented, with two additional functions added from original source code to add atoms. The first function was to gather all the free locations on top of the basement membrane and the second function adds the stem cells specified on the free locations available.

The pseudocode is shown in Algorithms 1 and 2:

#### Algorithm 1

Get list of free loactions

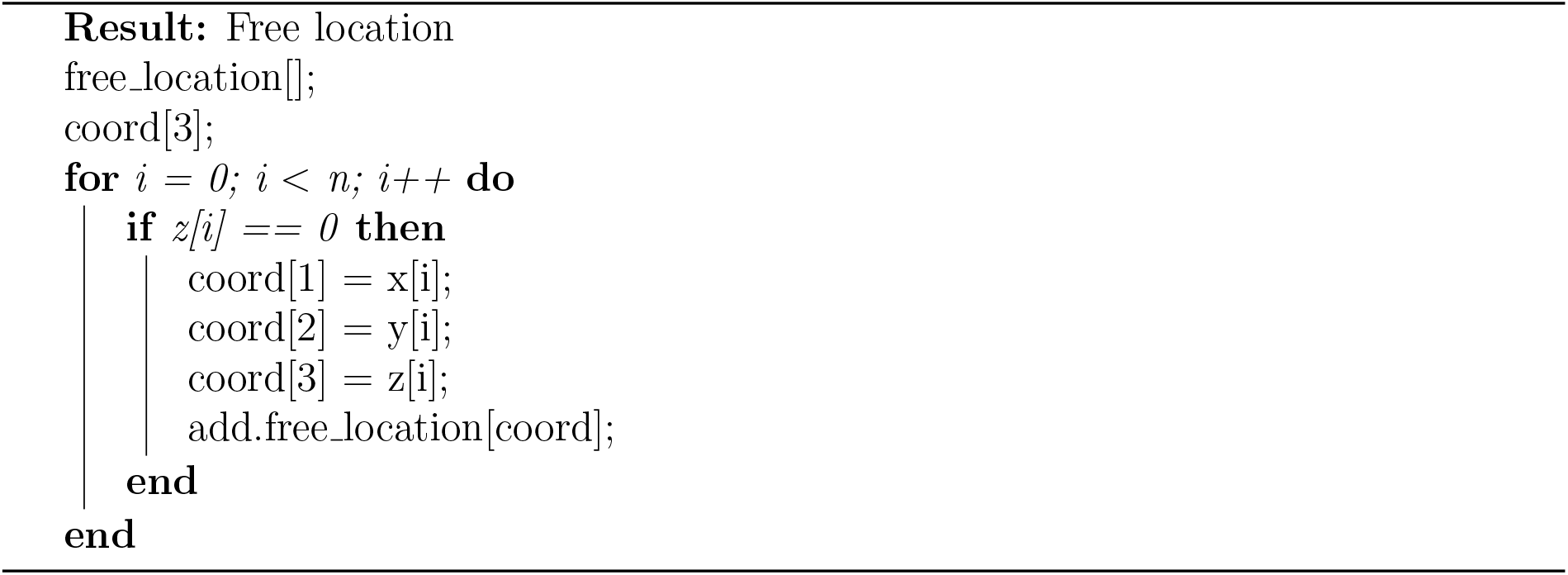

#### Algorithm 2

Initialising stem cell

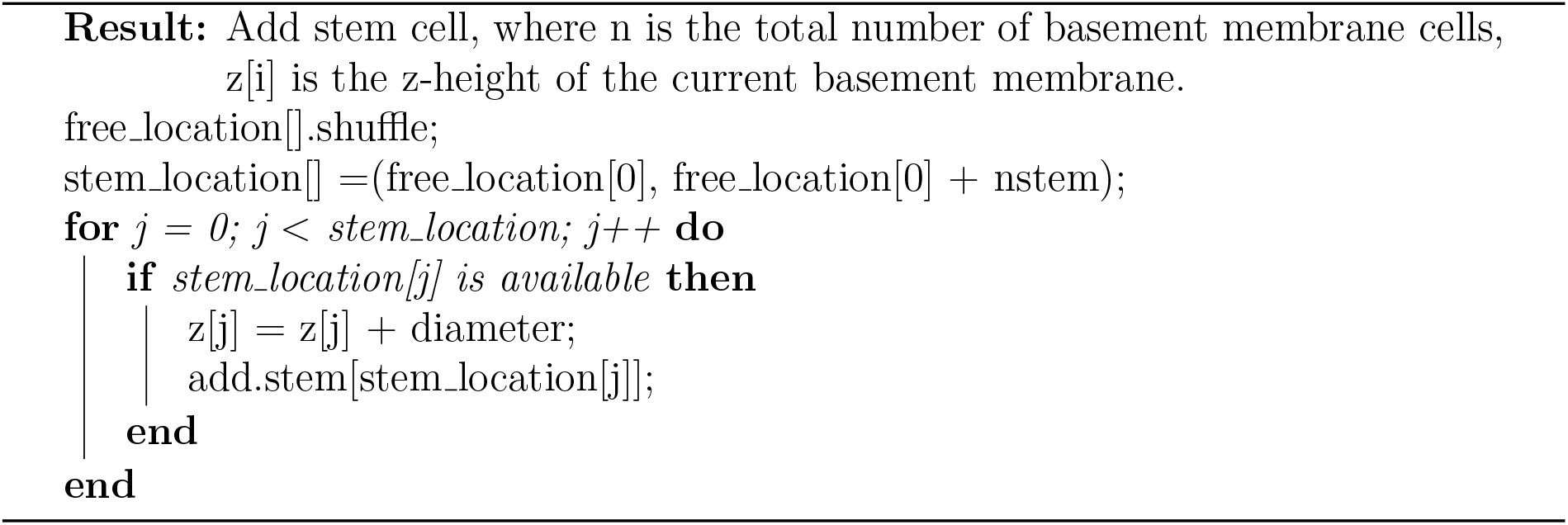

If the top of that cell is free, its coordinates are added to the list of free locations (*free_location*[]). Once the list of free locations is obtained, the list is shuffled and a new list is created of the same size of the number of stem cells to initialise ensuring that stem cells are randomly placed along the basement membrane surface.

### Mechanical relaxation

During cell division, overlapping cells may occur and so mechanical relaxation is required to update the cell’s position to prevent this. Mechanical relaxation is done by using a discrete element method and the Newtonian equations of motion. The movement of cell is done so by the following equation:

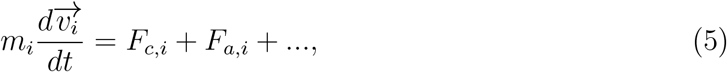

where *m*_*i*_ is the mass and 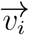 is the velocity. The types of forces acting on each cell are based on the contact and adhesive force.

The contact force *F*_*c,i*_ is a pairwise force exerted on cells to resolve the overlapping issues and to ensure that cells do not enter the basement membrane. This is based on Hooke’s law as follows:

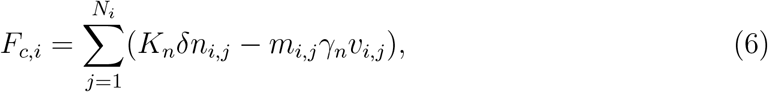

where *N*_*i*_ is the total number of neighbouring particles of *i, K*_*n*_ is the elastic constant for normal contact, *δn*_*i,j*_ are the overlapping distance between the center of particle, *i*, and neighbouring particle, *j, m*_*i,j*_ is the effective mass of particles *i* and *j, γ*_*n*_ is the viscoelastic damping constant for normal contact, and *v*_*i,j*_ is the relative velocity of the two particles.

The adhesive force between basal cells, stem and TA cells, *F*_*a,i*_ is the pairwise interaction modelled based on van der Waals force as follows:

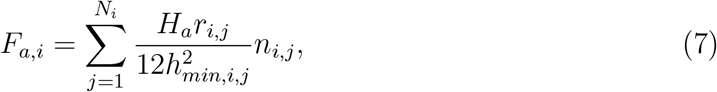

where *H*_*a*_ is the Hamaker coefficient, *r*_*i,j*_ is the effective outer-radius of particles *i* and *j*, and *h*_*min,i,j*_ is the minimum separation distance of the two particles, and *n*_*i,j*_ is the unit vector from particle *i* to *j*.

Table 1 summarises the mechanical interactions based on what type of cell it is interacting with.

**Table 1:** Summary of forces between each cell type and between the basement membrane in the model where the strongest force is 5+ and the weakest force with just 1+.

| Cell Type | Repulsive force | Adhesive force |
| --- | --- | --- |
| Same cell type | + | ++ |
| SC-TA | + | + |
| TA-D | + | + |
| SC-D | ++++ | + |
| SC-BM | + | +++++ |
| TA-BM | ++ | +++ |
| D-BM | +++++ | + |

### Spatial regulation for cell division in healthy epidermal development

Stem and TA cells are known to have different division directions such as horizontal or vertical. As stem cells reside only on top of the basement membrane, the division direction for daughter cells that are self cells will be horizontal from the parent cell’s position, while the daughter TA cell would be placed vertically (Figure 3a). In TA cells, the division direction will mainly be vertical to ensure that differentiated cells are on top. However, this concept requires additional spatial regulation as the surface of the basement membrane is not flat but a wave. Figure 3b shows how division would occur on the wave surface according to the horizontal and vertical rules. The stem cell on top first self proliferates and produces two stem daughter cells. However, in the next division, one of the stem cells divides asymmetrically and produces a daughter TA cell which takes the parent’s position while the daughter stem cell divides on top, thereby allowing stem cells to leave the stratum basale layer.

A solution to this is to include a check for stem and TA cells to be aware of their location, whether it is on top of the basement membrane or away from it. Figure 4 describes how different proliferative cell will divide.

**Figure 4:**
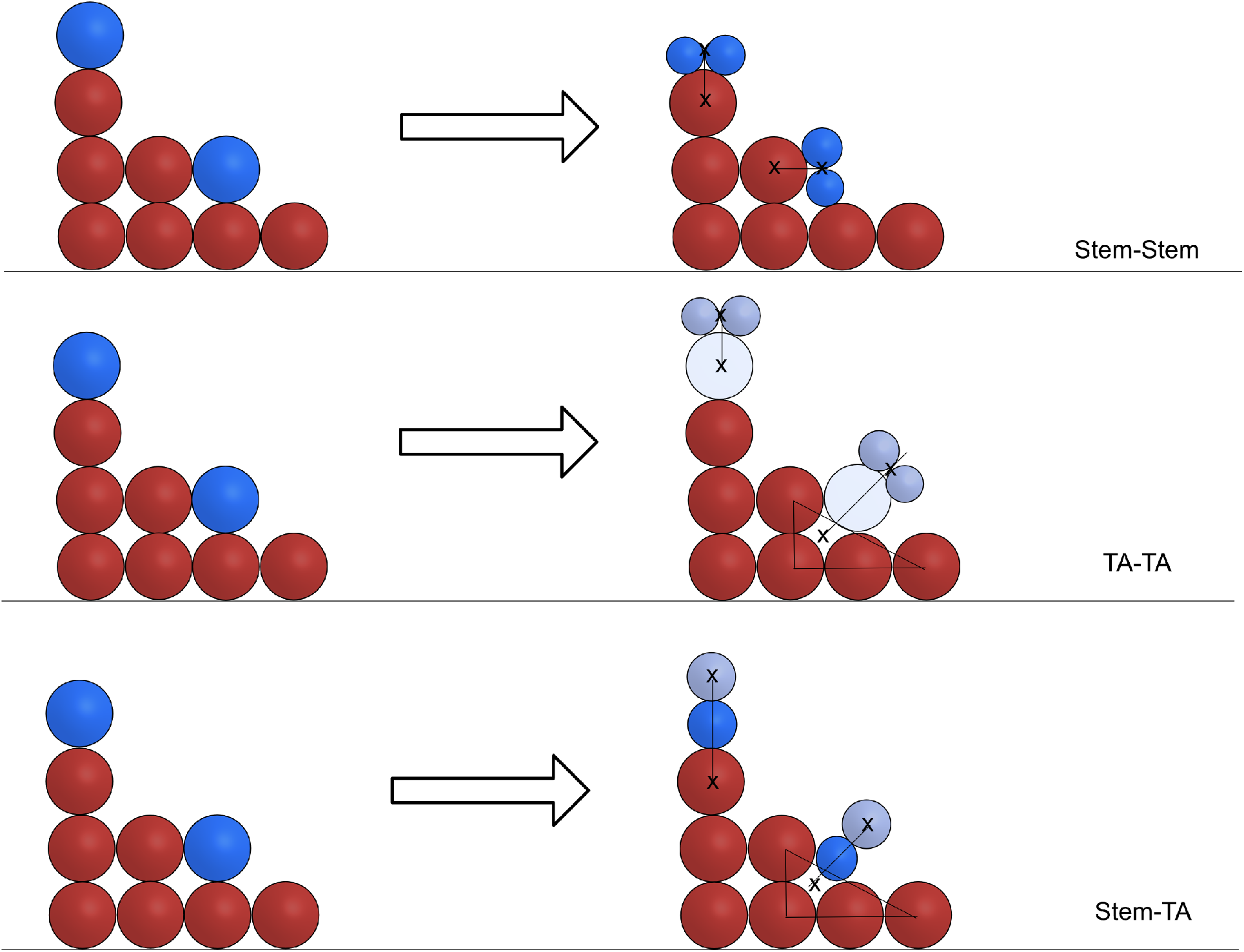
Schematic of spatial regulation implemented for stem cells. Self proliferation has been assumed to divide horizontally and TA daughter cells are assumed to be positioned vertically [19, 16, 20]. The concept of spatial regulation is based on these as follows: (Top) Stem cell self proliferation daughter cells are positioned side-by-side in the middle of the cell it is on top of. (Middle) Symmetric division behaves in a similar fashion when the division occurs on the top of the rete peg. If the cell was at the bottom of the rete peg, its position is calculated from the middle of the surrounding cells. (Bottom) Asymmetric division occurs in a similar fashion as symmetric, however, the daughter TA cell will always be on top of the daughter stem cell based on the assumption that TA cell will divide vertically.

### Chemical processes

Cytokine and calcium transport is described by using the diffusion-advection-reaction equation. In this section, we will briefly discuss the main concepts behind the chemical processes involved such as nutrient consumption and mass balance.

### Nutrient consumption

The rate of nutrient consumption is calculated within each voxel. The reaction rates are based on the Monod-based kinetics and can be defined as follows:

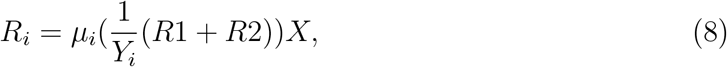

where *Y*_*i*_ is the yield coefficient, *R*1 is the growth during the normal state and *R*2 is the growth during psoriasis and *X* is the biomass density.

### Nutrient mass balance

Nutrient distribution within the simulation domain is calculated by solving a diffusion-advection-reaction equation (transport equation) for such nutrient as follows:

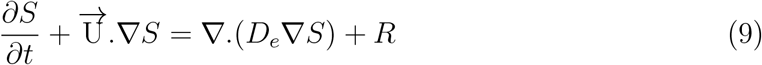

where the nutrient update, *R*, is calculated in the growth model, *S* is the nutrient involved. In this case, it is either one of the nutrients, calcium, GF, IL-22, or TNF-*α*.

The equation above is discretised on a Marker-And-Cell (MAC) uniform grid where the scalar *S*, is defined at the centre of voxel and velocity components U for x-, y- and z-axes are defined at the centre of the 6 faces of the voxel. The time derivative and spatial derivative are discretised by the Forward Euler and Central Finite Differences respectively.

### Transition to psoriasis

The biological and chemical processes are similar to the healthy epidermis model with some additions that will be further discussed in this section. The physical processes have been altered to take into account how the rete ridges deepen while hyperproliferation occurs. This is done to ensure that the wave-like shape is maintained as much as possible and to prevent the rete ridges from flattening out.

Psoriasis is modelled by activating an immune cytokine stimulus such as IL-22 and TNF*α* into the model, which causes an onset between 7 days to 14 weeks and a time lag between the time the patient gets an immunological response (such as from a streptococcal sore throat) to the appearance of the disease itself [7, 21, 22]. In this model, the immune cytokine stimuli is induced for a total of 7 days following the previously developed ODE model [10].

The growth rate of proliferative keratinocytes and differentiation in the psoriatic state increases by approximately three times in stem and TA in the model, which is similar to previous experimental data. In vivo experiments showed that the number of proliferating keratinocytes increases by 2-3 times to 6-8 times in psoriasis [23, 24]. This results in a 2-5 times increase in cell population [25, 26]. In this model, the increase in cell population, cell cycle and turnover times are predicted to be approximately 3-times than normal and are controlled by an “activator” switch that alters both the growth and division model.

### Alterations to epidermal structure

The transition from healthy and psoriatic epidermis results in a change to the epidermal structure causing the structure to be thicker and having deeper rete ridges.

Although there have been studies on identifying how the rete ridges form in the epidermis, the main cause it still unknown. In a study [27] on rete ridges in the oral cavity, the results showed that rete ridges form due to the sucking action in the mouth as a baby. This sucking action along with a higher density of stem cells causes a change in the oral cavity and forms the rete ridges. The depth of rete ridges is determined by the density of stem cells. Hence, the higher the stem cell density, the deeper the rete ridges. A similar mechanism of action can be assumed in the case of the skin, where physical forces come to play, causing rete ridges to form in the epidermis.

The model assumes a similar mechanism where mechanical forces are used to mimick this “sucking” action causing the deepening of rete ridges during psoriasis. To model these changes, the following changes have been made when switching states:

- Unfreeze the basement membrane to allow the deepening of rete ridges.
- Additional forces applied to stem and TA cells to ensure that these cells remains in the lower epidermal layers as the rete ridges start to deepen (Figure 5).
- An increase in height of the domain box to simulate a thicker epidermis.

**Figure 5:**
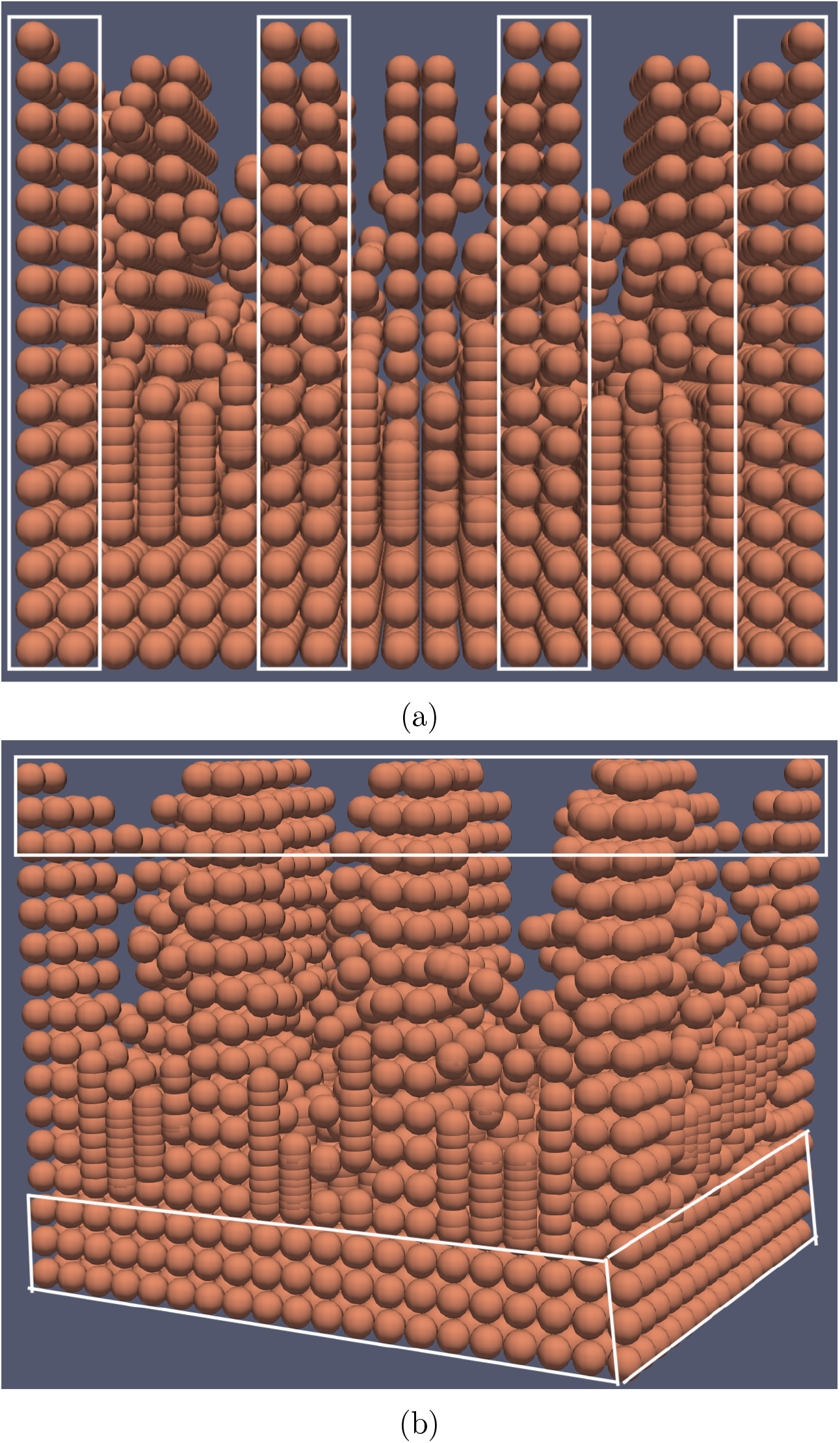
Regions highlighted in the white boxes where basement membrane (in orange) are “frozen” to ensure that wave shape is maintained when transitioning to psoriasis. Peak columns are frozen to ensure that only the toughs of the wave moves. Hence, producing the deepening of rete pegs are more cells are produced in the epidermis. (b) The top and bottom of the rete pegs are “frozen” to ensure the wave shape is maintained. The top of the rete peg also has an additional adhesive force between stem and TA. cells during the transitional state. This ensure that stem cells remain on top of the basement membrane and for TA cells to remain in the lower layers of the epidermis.

In addition, to ensure that the wave shape is maintained in the model, the basement membrane has been sub-divided into different regions where the peaks and toughs of the waves are sub-divided into columns. The column regions with the peaks are “frozen” in place to ensure that only the toughs are deepened, hence, producing deeper rete ridges.

### Alterations to TA cell division

The transition to psoriasis differs slightly as the model does not start off with a clean slate of just stem cells but rather a domain filled with the various cell types. In addition to the difference in starting state, TA cells may also reside in the stratum basale layer which lies in the rete ridges. Hence, additional rules applied to TA cells are required to ensure that 1. TA cells do not push sideways causing the epidermal structure to lose that wave-like shape during the transition to psoriasis, 2. epidermal layer stratification is maintained. The changes made to the direction of how TA cells divide ensured that they divide vertically [28, 29], or oblique [30] manner as seen in previous models and clinical studies. Figure 6 shows a time-lapse immunofluorescence images tracking the direction of proliferative cell division in mice [30]. The study investigated in different parts of hairless mice such as the dorsal, ear, hindpaw and tail skin. It was revealed that most division along the basement membrane in the dorsal and ear epidermis were parallel, whereas in the hindpaw and tail epidermis, cell division occurs in parallel and oblique in direction. Although, the area investigated differs slightly from how the human epidermal structure is, this provides some insights on how TA cell division may differ slightly when the epidermis has been fully developed. In this model, we have adapted this concept and applied it to TA cell division in the development of psoriasis.

**Figure 6:**
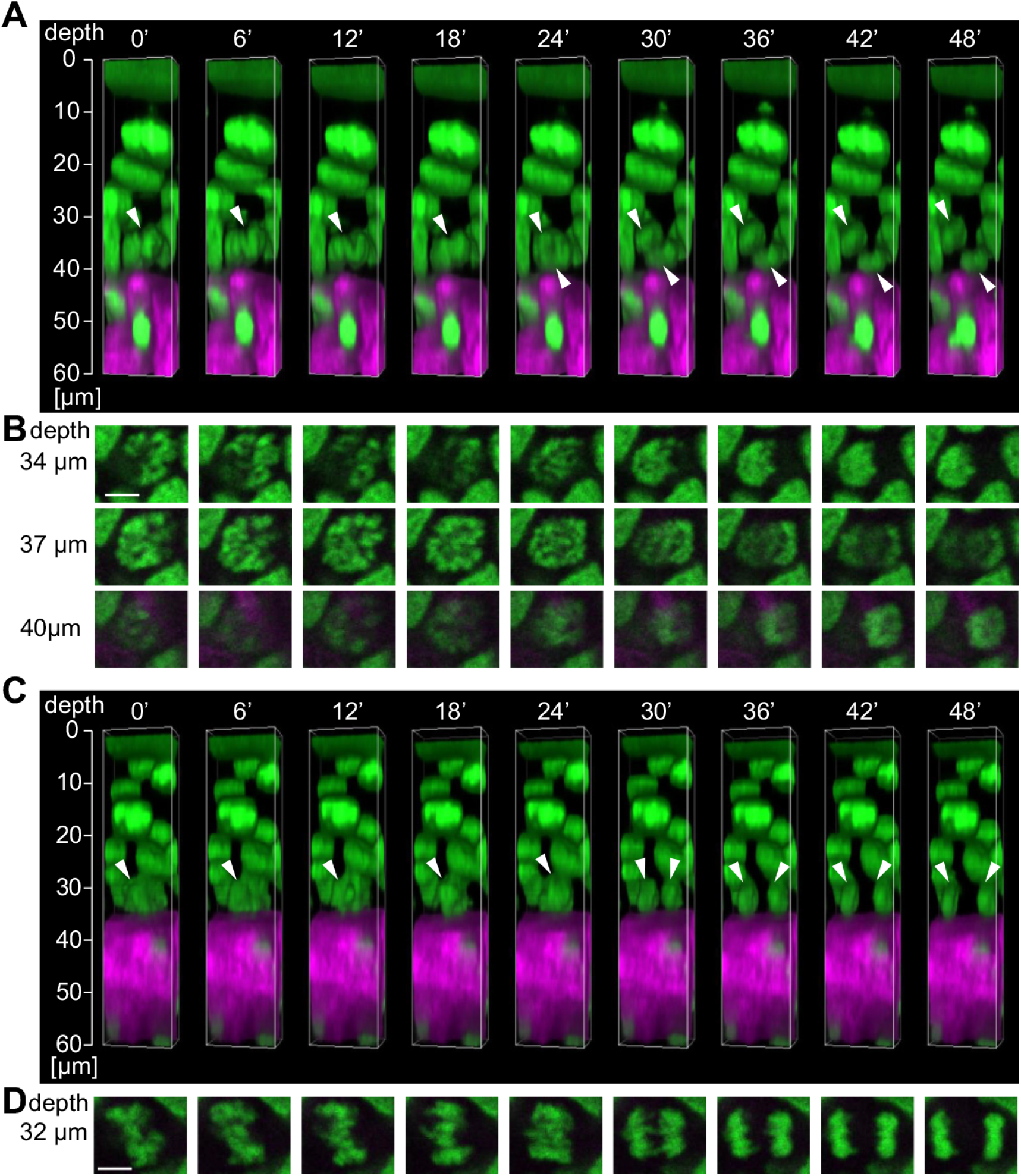
Figure adapted from [30]. A four-dimensional imaging of cell division in the hind paw epidermis in a living mouse. (A) A reconstruction 3D image of oblique division to the basement membrane. Images are take every 6 minutes. (B) Time-lapse x- and y-axis images of mitotic cells in (A) at different depths. (C) Reconstructed 3D images of parallel division along the basement membrane. Images are take every 6 minutes. (D) Time-lapse x- and y-axis images of mitotic cells in (C) at depth of 32*µ*m from the skin surface. White arrows represents the cell dividing. Scale bar = 5*µ*m.

### Introduction of Immune Cytokine Stimulus

In addition to calcium and endogenous GF involved in the growth and proliferation of stem and TA cells, a single immune cytokine stimulus is initialised in the model to induce psoriasis. The cytokine stimulus initialised is maintained by introducing T cells into the system, where they reside in the basement membrane and the dermis region. The function of T cells has been simplified where they do not grow and migrate out of the basement membrane. The introduction of T cells is there to mainly regulate the amount of stimulus in the system, to activate and maintain psoriasis.

### Parameters

The parameters can be found in Table 2. The division probabilities for TA cell (*P*_*c*_ and *P*_*d*_) are based on the expanding and balanced tree division probabilities found in [12] where they have labelled and measured the types of division that occur. The division probabilities was only used for TA cells in the normal epidermal developed as in the paper, results obtained were based on the proliferative properties of a cell. Therefore, it will not be a good representative for stem cells as in other studies, it has been said that only about 5% of stem cells are cycling. Hence, we assumed that the cells that were considered to have proliferative properties to be TA cells. Parameter estimation was done for the division probabilities for stem cells with the assumption that *P*_0_ in the balanced state was 0.5 to maintain a steady state. It was found that for the model to maintain a steady state and obtain the cell population ratio that was within the normal range, 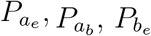 and 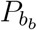 values were 0.05, 0.05, 0.1, 0.7 respectively. While in psoriasis, 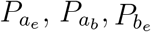 and 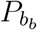 values were 0.05, 0.05, 0.4, 0.4 for stem cells, respectively, and 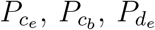 and 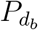 values were 0.25, 0.1, 0.25 and 0.05 for TA cells, respectively.

**Table 2:** Model parameters

| Normal epidermis |  |  |  |  |
| --- | --- | --- | --- | --- |
| Parameter description | Parameter | Value | Units | Reference |
| Stem cells initialised | $SC_{init}$ | 290 | Cells | Estimated |
| SC maximum growth rate | SC $\mu_{max}$ | 1.28e-06 | $s^{-1}$ | Modified from [7] |
| TA maximum growth rate | TA $\mu_{max}$ | 3.21e-06 | $s^{-1}$ | Modified from [7] |
| Ks value for growth factor | $Ks_{gf}$ | 2.5e-7 | $kgm^3$ | Estimated |
| Ks value for calcium | $Ks_{ca}$ | 2.5e-3 | $kgm^3$ | Estimated |
| SC symmetric division in expanding stage | $Pa_e$ | 0.05 | - | Estimated |
| SC symmetric division in balanced stage | $Pa_b$ | 0.05 | - | Estimated |
| SC asymmetric division in expanding stage | $Pb_e$ | 0.1 | - | Estimated |
| SC asymmetric division in balanced stage | $Pb_b$ | 0.7 | - | Estimated |
| TA symmetric division in expanding stage | $Pc_e$ | 0.02 | - | [12] |
| TA symmetric division in balanced stage | $Pc_b$ | 0.32 | - | [12] |
| TA asymmetric division in expanding stage | $Pd_e$ | 0.1 | - | [12] |
| TA asymmetric division in balanced stage | $Pd_b$ | 0.18 | - | [12] |
| Maximum TA division | $max_{TA}$ | 4 | - | [7] |
| Diffusion coefficient | $Ks$ | 1.0e-9 | $m^2s^{-1}$ | [31] |
| Extracellular Calcium initialised | $eCa^{2+}$ | 1.0e-7 | $kgm^3$ | Estimated |
| Psoriatic epidermis |  |  |  |  |
| SC symmetric division in expanding stage | $Pa_e$ | 0.05 | - | Estimated |
| SC symmetric division in balanced stage | $Pa_b$ | 0.05 | - | Estimated |
| SC asymmetric division in expanding stage | $Pb_e$ | 0.4 | - | Estimated |
| SC asymmetric division in balanced stage | $Pb_b$ | 0.4 | - | Estimated |
| TA symmetric division in expanding stage | $Pc_e$ | 0.25 | - | Estimated |
| TA symmetric division in balanced stage | $Pc_b$ | 0.1 | - | Estimated |
| TA asymmetric division in expanding stage | $Pd_e$ | 0.25 | - | Estimated |
| TA asymmetric division in balanced stage | $Pd_b$ | 0.05 | - | Estimated |
| Maximum TA division | $max_{TA}$ | 5 | - | [7] |
| Diffusion coefficient | $Ks$ | 1.0e-9 | $m^2s^{-1}$ | [31] |
| Immune cytokine stimulus initialised | $Stim$ | 5.0e-4 | $kgm^3$ | Estimated |
| Ks value for stimulus | $Ks_{stim}$ | 2.e-7 | | Estimated |

## Results

### Rete Peg Formation and Stem Cell Initialisation

The rete peg formation was based on the sine wave formation. Stem cells initialisation are based on the new fix command implemented which stores a list of the available locations on top of the basement membrane and randomly adds the stem cells on top of it by shuffling the free locations list and having a new list with the available locations the same size as the number of stem cells initialised. Figure 7 shows the basement membrane (in red) and stem cells (in blue) at the start of the simulation. The rete pegs in the normal epidermis will remain in place as cell population increases as seen in Figure 9a.

**Figure 7:**
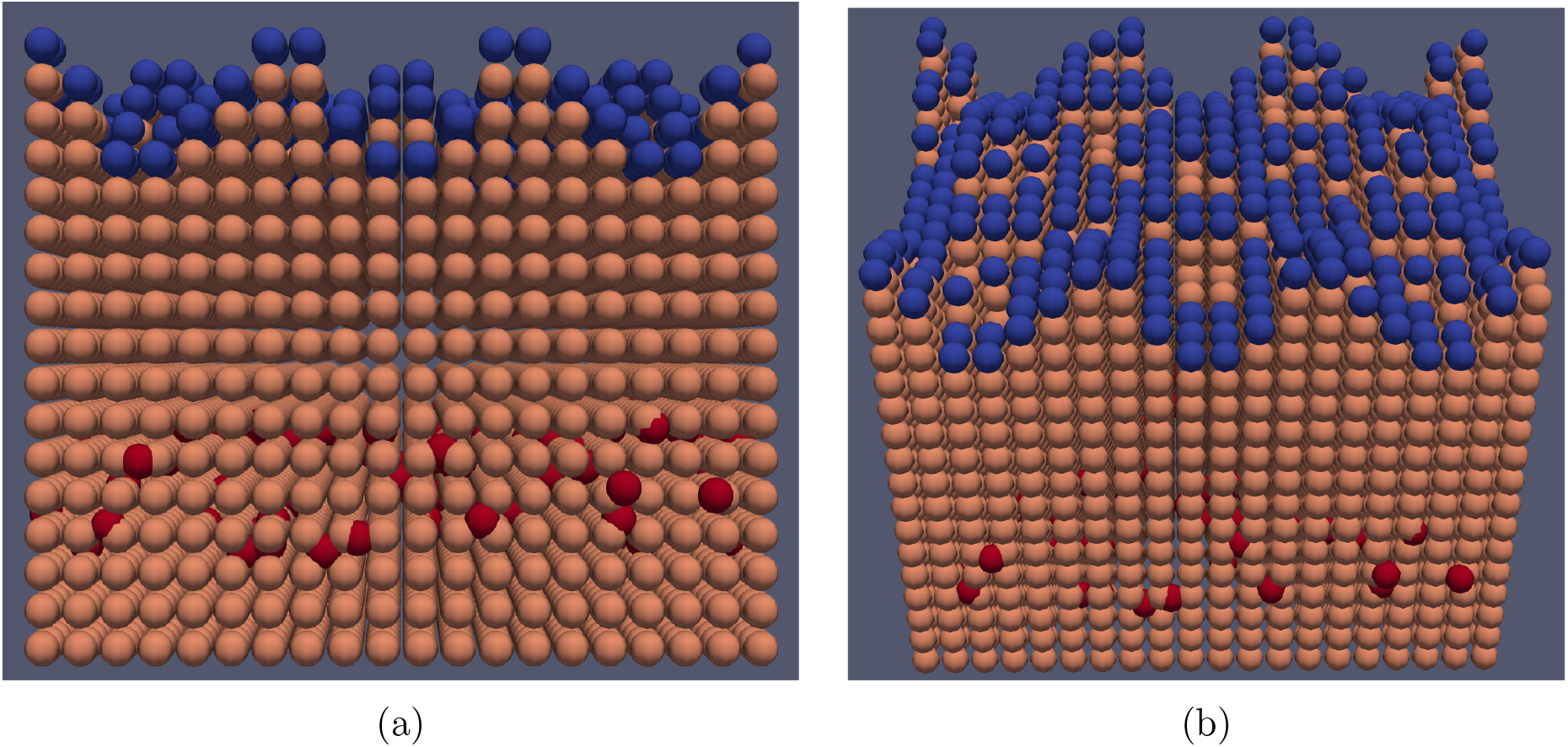
Screenshot of the wave formation use in the model. The basement membrane (in orange) is formed by creating atoms using the LAMMPS command “*create*_*atoms* and the structure is fixed to ensure that no movement occurs during epidermal growth. Stem cells (in blue) are initialised on top of the basement membrane with free locations available.

**Figure 8:**
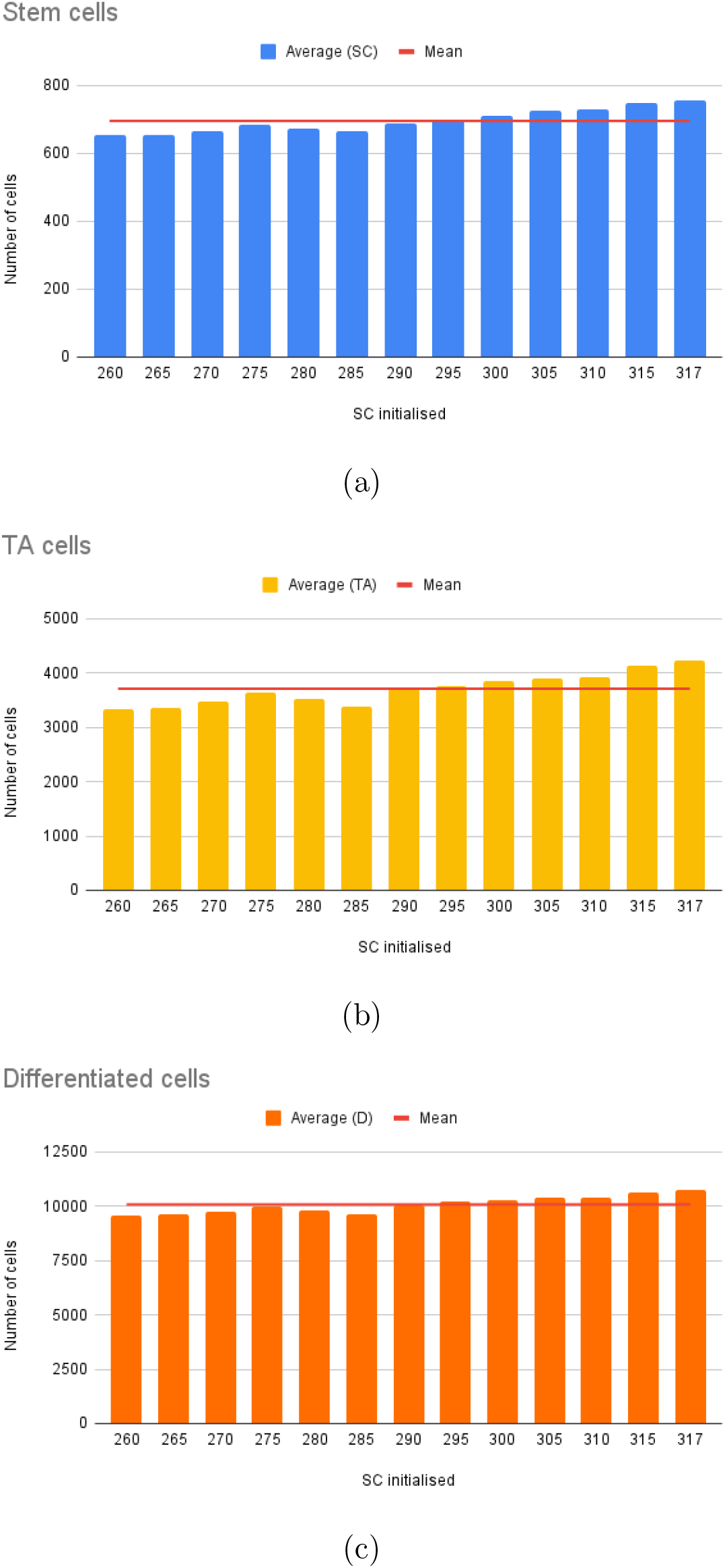
Average cell population based on the number of stem cells initialised where the best range was found to be between 260 to 317 stem cells initialised. (a) Stem cell population in steady state. (b) TA cell population in steady state. (c) Differentiated cell population in steady state. The average out of all simulations are represented by the horizontal line in the plots, where the average numbers are 695, 3709, and 10080 for stem, TA and differentiated cells respectively. The results obtained were measured based on an average run of 10 simulations for each test case. Table **??** summarises the parameters used.

**Figure 9:**
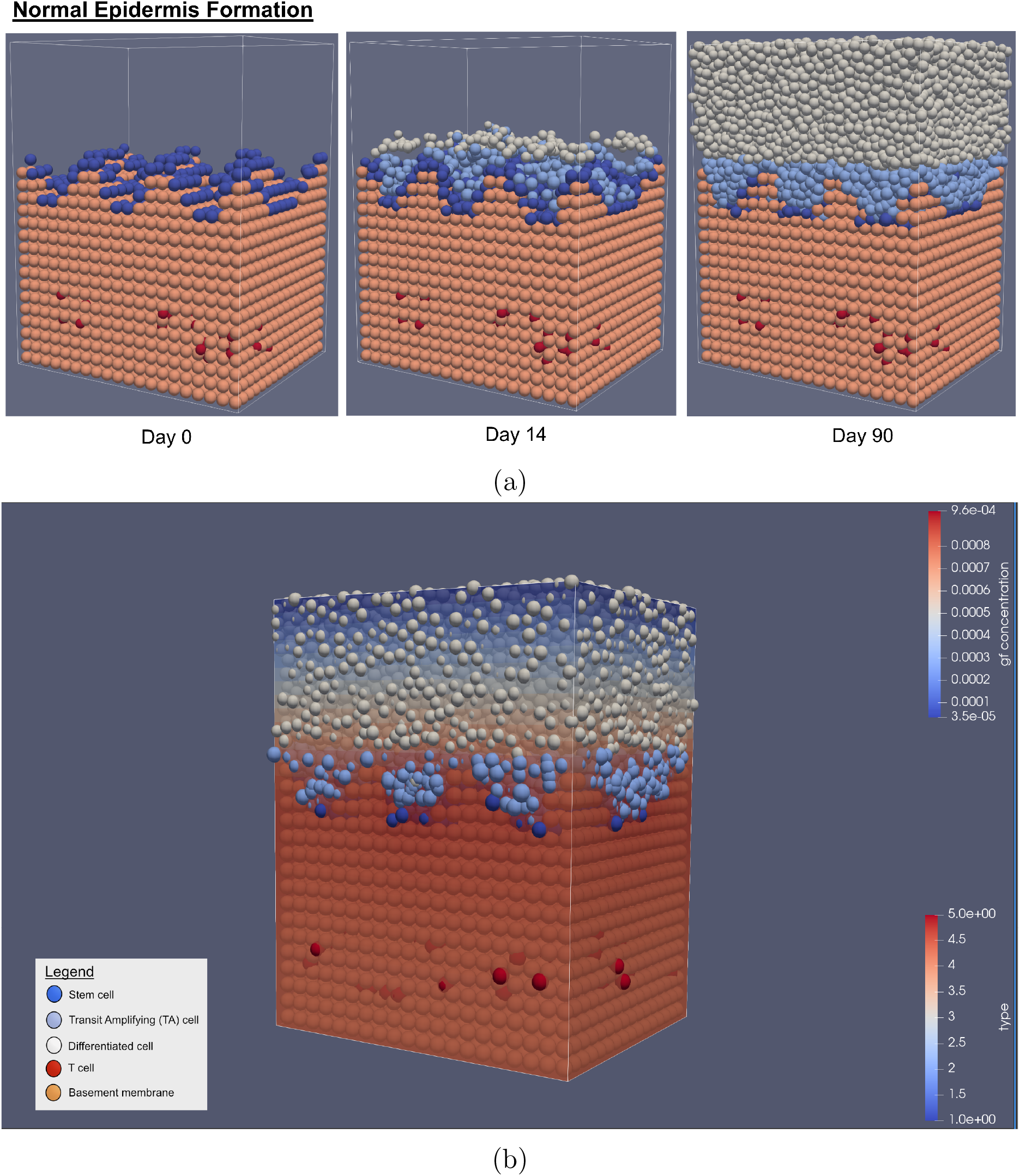
Visualisation output from the model during healthy epidermal formation. (a) Snapshot of the model at three different time points (Day 0, 6 and 30). (b) Snapshot of diffusion gradient in the model. In this case, growth factor, with the highest concentration being around the stratum basale layer where stem and most TA cells reside. Cells represented: stem cells (in dark blue), TA cells (in light blue), differentiated cells (in white), T cells (in red) and basement membrane (in orange).

**Figure 10:**
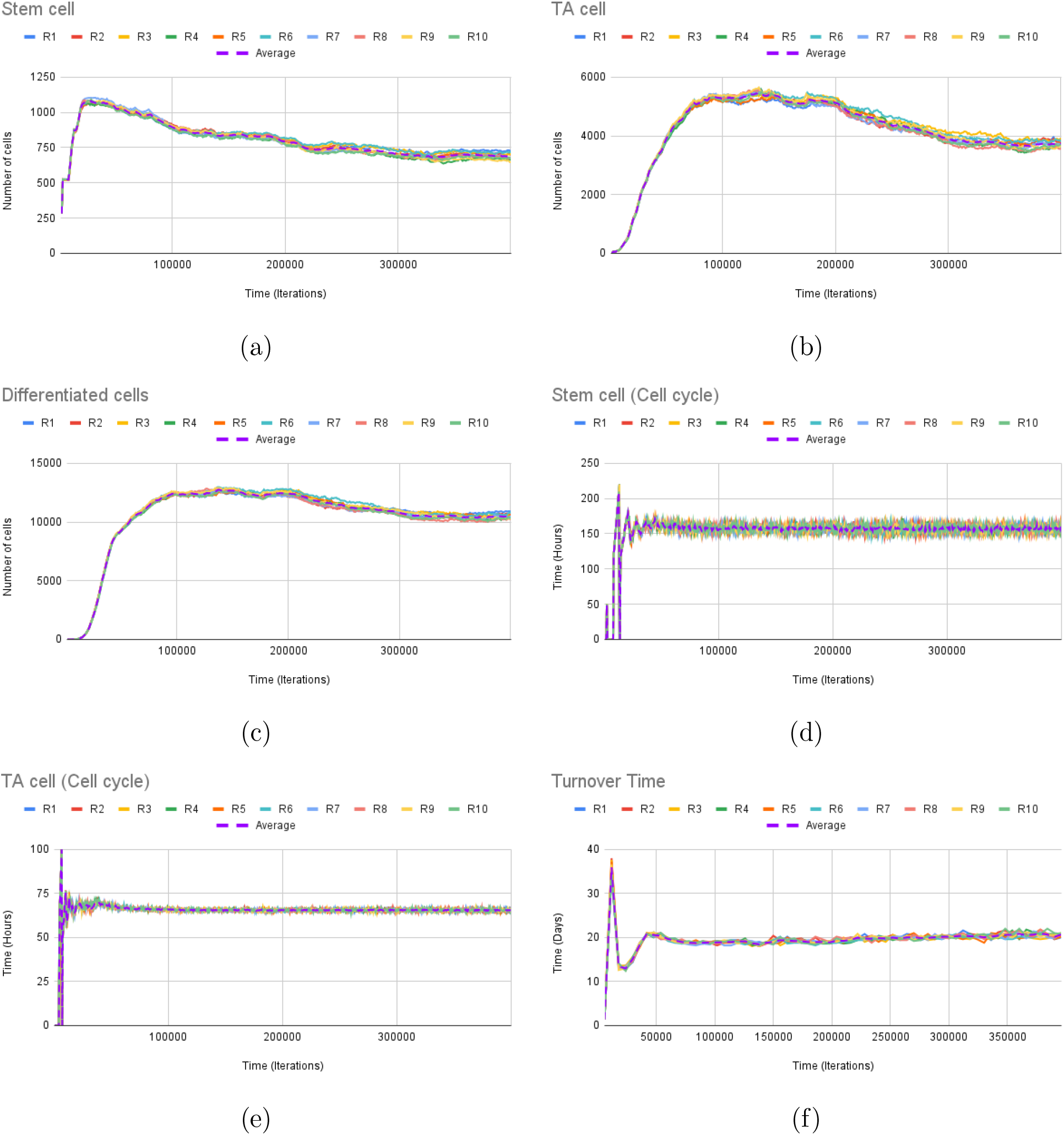
(a-c)Cell numbers plot of stem cells (a), TA cells (b) and differentiated cells (C) in healthy epidermis. The average ratio of stem, TA and differentiated cells are 4.49% (693), 24.3% (3751) and 67.98% (10 494), respectively. (d-e) Cell cycle for stem (d) and TA cells (e). The average cell cycle times predicted are 157.2 and 65.5 hours, respectively. (f) Total turnover times for both proliferative and differentiated compartment. The average turnover time in the model is 20.5 days in the steady state. The simulations were ran 10 times under different random seed (R1-10) with the average plotted in purple dotted line. Each simulation ran under the same initial conditions such as the extracellular calcium diffused, growth rates, and diffusion rates. Table 2 summarises the parameters used.

The number of stem cells initialised in the system has been tested from a range of 100 to 317 cells at intervals of +10 cells initially and further refined to +5 cells step. The maximum number of stem cells that can be initialised to cover the entire surface of the basement membrane is 317. If more than 317 stem cells are initialised, they are “removed” from the system as there is insufficient space for them to be placed on top of the basement membrane. The results showed that for the ideal cell densities to populate the epidermis, the number of stem cells required for initialisation is found to be between 260 to 317 cells. If less than 260 stem cells are initialised, the model fails to reach the ideal cell densities and proportions.

In addition, stem cells are randomly initialised on the basement membrane surface and no specific location is set. Previous studies on stem cell location report conflicting results which show stem cells either residing at the bottom or top of the rete pegs. In this work, the location of stem cells was randomly initialised due to two main reasons. Firstly, the model starts off with the basement membrane shaped as a wave structure, which means that the stem cell growth and division did not form this irregular shape as compared to embryonic studies which looked at how the epidermis was formed from the beginning of time. Secondly, this model does not study how the epidermis forms from an embryo. Therefore, the number of stem cells has to start at a larger number to obtain a well and ideally proportioned epidermis with all three types of cells as stem cell growth is slower than TA cells.

### Cell Population Density

The cell population density of healthy epidermis reported in previous studies was of 73,952 ± 19,426 *cells/mm*^2^ (SD) [32] with each cell population ratio estimated at 4.7%, 26.4% and 40-66% of them being stem, TA and differentiated cells respectively with keratinocytes comprising of 96% of all epidermal cells [7]. As the model only takes keratinocytes into account, the model has a cell population ratio of 4.49% stem cells, 24.3% TA cells and 67.98% differentiated cells, which translates to the cell population ratio of keratinocytes being 4.3%, 23.33% and 65.2%.

A previous computational study [10] showed that the cell population ratio remained the same in both healthy and psoriasis. In addition, the overall number of epidermal cells is known to increase by up to two to five times as compared to the healthy epidermis [25, 33]. The model predicted similar cell populations with the ratio being 4.51% (2,103), 25.7% (11,976) and 68.8% (32,088) for stem, TA and differentiated cells, respectively. As only keratinocytes are taken into account, this translates to a cell population ratio of 4.33%, 24.67% and 66% for stem, TA and differentiated cells. This is similar to the cell population ratio in the healthy epidermis with 4.3%, 23.33% and 65.2% being stem, TA and differentiated, respectively. The overall number of epidermal cells has also increased by approximately three times as seen in Figure 11 showing the model outputs and average for 10 simulation runs under different random seeds.

**Figure 11:**
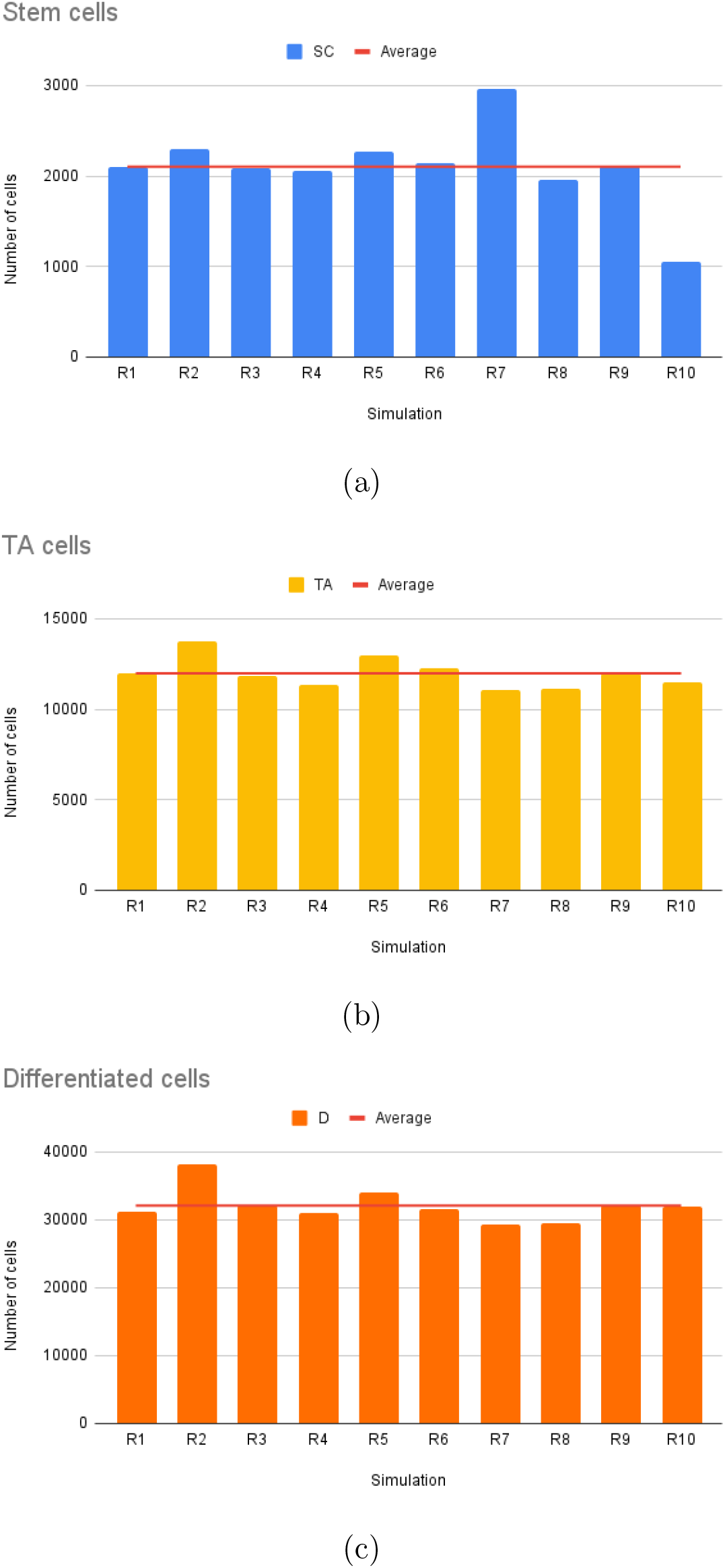
Average cell population numbers in the psoriatic steady state. (a) Stem cell population in steady state. (b) TA cell population in steady state. (c) Differentiated cell population in steady state. The average out of all simulations are represented by the horizontal line in the plots, where the average numbers are 2,103, 11, 976, and 32, 088 for stem, TA and differentiated cells respectively. The overall number of epidermal cells has increased by 3-times as compared to the healthy epidermis. The results obtained were measured based on an average run of 10 simulations for each test case. Table 2 summarises the parameters used.

**Figure 12:**
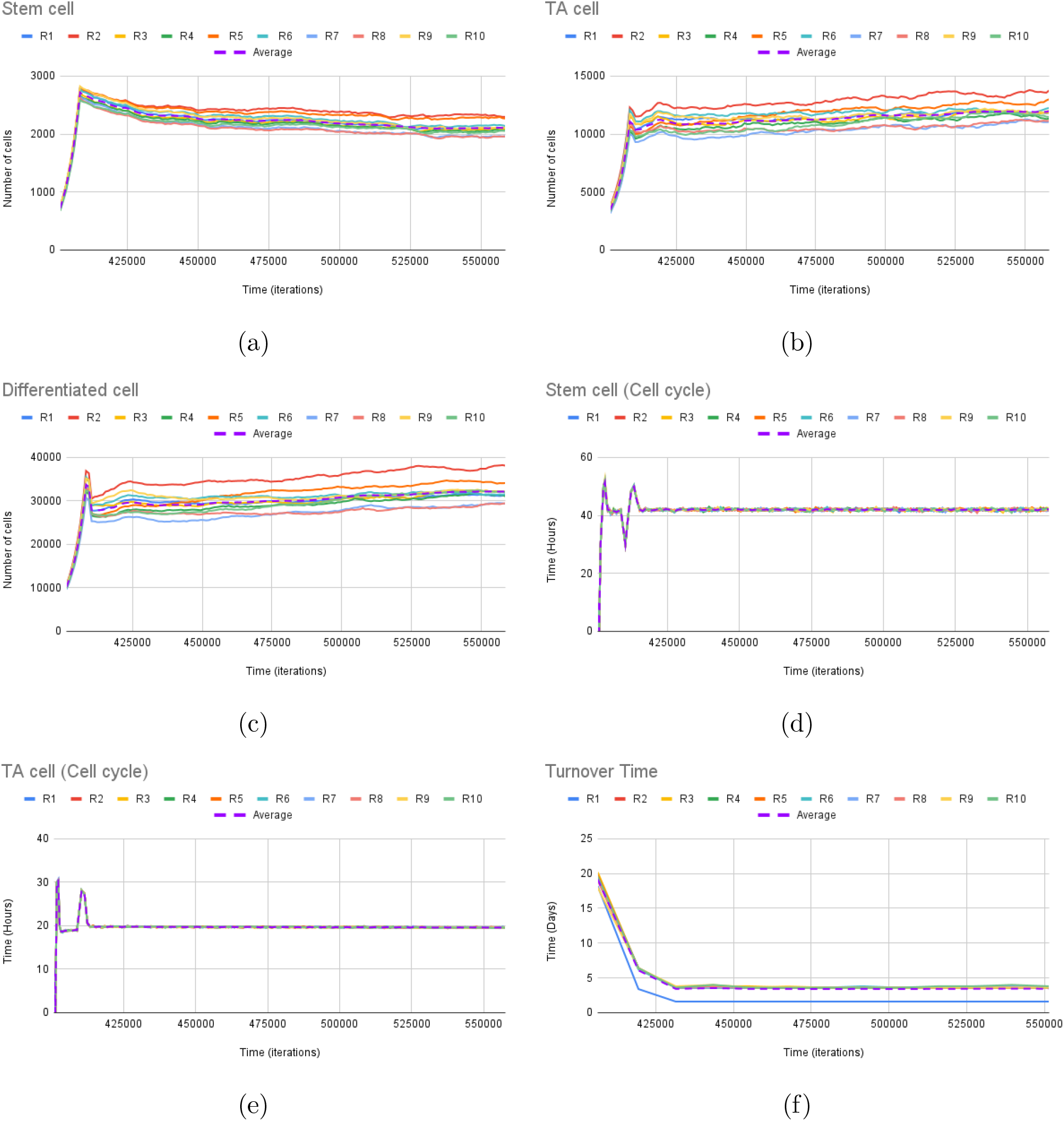
(a-c)Cell numbers plot of stem cells (a), TA cells (b) and differentiated cells (c) in psoriasis. The average ratio of stem, TA and differentiated cells are 4.51% (2103), 25.7% (11 976) and 68.8% (32 088), respectively. (d-e) Cell cycle for stem (d) and TA cells (e). The average cell cycle times predicted are 42 and 19.7 hours, respectively. (f) Total turnover times for both proliferative and differentiated compartment. The average turnover time in the model is approximately 4 days in the steady state. Overall, the model is able to simulate psoriasis with the cell cycle and turnover times approximately 3.5- and 4-times faster, respectively. The simulations were ran 10 times under different random seed (R1-10) with the average plotted in purple dotted line. Each simulation ran under the same initial conditions such as the immune cytokine stimulus initialised, growth rates, and diffusion rates. Table 2 summarises the parameters used.

### Turnover and cell cycle times

The reported cell cycle times in clinical studies of healthy epidermis were found to be between 100-200 hours [34, 7] and 50-65 [32, 7] hours in stem and TA cells respectively. The measured cell cycle time range in the model were approximately 152.9 to 162.0 hours for stem cells and 65.0 to 66.0 hours for TA cells with an average of 157.2 and 65 hours, respectively.

The turnover time is calculated based on [35, 36] and [37] where the total epidermal turnover time is the sum of the proliferative and differentiated compartment. Each compartment’s turnover time is based on Equation 10. Hence, giving us Equation 11.

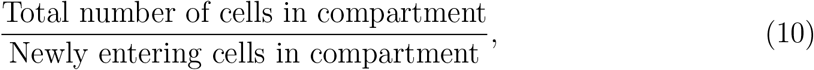

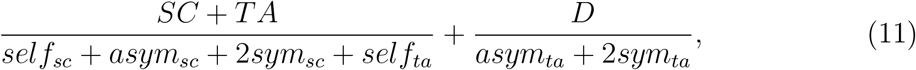

where the first term is for the proliferative compartment, stem and TA cells, and the second term is for the differentiated compartment, differentiated cells. Variables *self*_*x*_, *asym*_*x*_ and 2*sym*_*x*_ represent the three types of division modelled, self-proliferation, asymmetric and symmetric division while _*x*_ represents the cell undergoing division, which could either be a stem or TA cell. It is also important to note that symmetric division produces two daughter cells of a different type from the mother cell, hence, producing two cells for that compartment. For example, 2*sym*_*sc*_ will produce two TA cells from a stem cell, in the proliferative compartment.

In this work, the stratum corneum has not modelled and only takes into account the proliferative and differentiated compartments. Therefore, the model predicts the total turnover times to be between 19.6 and 21.3 days with an average of 20.5 days, slightly lower than the measured 23-25 days in [25].

In psoriasis, it is known that the cell cycle and the turnover times are three to five times faster than in healthy epidermis [25, 36, 33]. The predicted cell cycle and turnover times in psoriasis are approximately 4-times shorter. The average cell cycle times are 47 and 20 hours for stem and TA cells, respectively, while the total turnover time was 5-times faster at approximately 4 days.

## Discussion

Psoriasis is a chronic inflammatory skin disease which affects the patient’s quality of life and currently has no cure. Computational models provide not only scientists but medical practitioners a better understanding of the disease formation, with the aim of better understanding the disease formation and providing specific treatment options. A previously developed 2D model of psoriasis [7] and NB-UVB clearance looked into solving some of these issues when understanding NB-UVB phototherapy treatments. However, the 2D model has some limitations such as the lack of cell-cell interactions, spatial considerations, and high NB-UVB dose given for each treatment. These do not depict how the epidermis develops and behaves and the actual treatment dose given in clinic. This work focused on solving some of these limitations with the aim to develop a 3D computational model of immune cytokine interaction and epidermal homeostasis in psoriasis.

### Limitations

The 3D model developed is a simplification of how the epidermis develops in reality. It takes into account the three main types of keratinocytes involved in the epidermis and their interactions with a single immune cytokine stimulus to progress to psoriasis. However, in reality, there are more than the three types of cells mentioned such as granular cells and corneocytes [38, 39, 28] and different, multiple immune cyotkine stimulus and T cell species [40, 41, 42] that may be involved in the development of psoriasis. The limitations are listed as follows:

- The model starts as an embryonic state which may have a different initial rate of growth and transition to psoriasis.
- The model has been simplified to include only the main three types of keratinocytes - stem, TA and differentiated cells. However, in reality, differentiated cells can be further subdivided and labelled according to where it is located in the epidermis. For example, TA cells will differentiate to growth arrest cells which will differentiate to a spinous cell in the stratum spinosum layer. The cell will go on to differentiate further into a granular cell in the stratum granulosum and corneocyte stratum corneum layer [38, 39, 28].
- Corneocytes are differentiated cell found in the upper most layer of the epidermis, the stratum corneum. Corneocytes are not spherical in shape but a flat cells which has excreted out its contents in preparation to be desquamated out of the epidermis. In this model, corneocytes have not been taken into account and the action of differentiated cells flattening is not modelled. Therefore, the turnover time in the model is slightly shorter than what is found in literature as shedding does not occur.
- The model does not include other cells that are present in the epidermis such as hair follicles and melanocytes. Hair follicles produce the hair on our skin [13], and melanocytes produce melanin (the pigment responsible for our skin colour) [43, 44, 45, 46].
- A single immune cytokine and T cell species are modelled. However, in reality, there are many different types of T cell and cytokine species which are known to cause psoriasis such as IL-22, IL-23, IL-17, TNF*α* and so on [1]. Each cytokine is known to affect a different inflammatory response that can cause different severity of the disease. Hence, they are important in biological treatment as they target specific cytokines [1, 42].

### Future Work

The 3D computational model simulates how a healthy epidermis forms and its transition to psoriasis using a single immune cytokine stimulus. The model provides the first step as a baseline model to dive further into how we can use computational models to better understand the disease and to model various treatment options such as NB-UVB phototherapy or biological treatments. Some proposals for future studies include modelling NB-UVB phototherapy treatments, exploring different doses and frequencies of NB-UVB treatments and using single or combination mechanisms of action for clearance, similar to the 2D model previously developed by [7]. In addition, the model outputs can also be fed into a machine learning algorithm previously developed [10] to make predictions of how each simulation can be clustered to emulate real life patients.

It can also be modified to investigate specific immune cytokine stimuli and T cell species or a combination of them to better understand how each cytokine affects the skin to cause different variations of psoriasis. This would allow us to explore how different biological treatments aid clearance of psoriasis by targeting specific cytokines or may be used in combination therapies.

Apart from modelling psoriasis treatments, some potential work include tackling some of the limitations mentioned in the current 3D model. This includes modelling the upper epidermal layers where differentiated cells flatten out and eventually shed off to develop a model where all layers of the epidermis are included which can be used to explore other skin diseases apart from psoriasis. Next, the inclusion of other cell types such as melanocytes as they are known to affect UVB phototherapy treatments [43, 44, 47, 48, 45]. Melanocytes are a type of dendritic cells that produce melanin, the pigment responsible for skin colour, located in the stratum basale layer. Melanin is known to absorb UV, preventing DNA damage to keratinocytes. The amount of melanin in the body correlates to how much UV protection the skin gets, hence, patients of colour has a lower susceptibility of UV damage as it offers more protection [49]. It has been suggested that phototherapy is effective in patients of colour and may require a higher dose [50, 51]. A disadvantage of phototherapy is the risk of post-inflammatory hyperpigmentation (i.e. darkening of the skin), which occurs due to exposure to NB-UVB phototherapy treatments, and may not be acceptable to all patients [52]. It will be interesting to investigate how different NB-UVB doses affect patients of colour and how we may use computational modelling to predict the minimal dose required.

In conclusion, the 3D computational model present insights on how the epidermis forms and how immune cytokines trigger psoriasis, altering not only the structure of the epidermis but also alters how proliferative cells, stem and TA, divide. Directions for future work include modelling psoriasis NB-UVB and biological treatments but also tackling some of the limitations in the current model so that it can be used hand-in-hand or as a predictive tool for future clinical studies as well.

## Code and Data

The code can be found on https://github.com/dinikap/NUFEB.git, under the branch “psoriasis-remote”. The source code for the psoriasis package can be found in “src/USERPSORIASIS”. The working examples can be found in “examples/epidermis-test” and “examples/psoriasis-test” for the normal and psoriatic epidermis, respectively. In “examples/epidermis-test/epidermis-1”, runs the normal epidermal formation and in “examples/psoriasis-test/psoriasis-1” runs the transitional state to psoriasis using random seed 10. For the steady state model for the same random seed, run “examples/psoriasis-test-psoriasis-10”.

